# Climate and developmental plasticity: interannual variability in grapevine leaf morphology

**DOI:** 10.1101/030957

**Authors:** Daniel H. Chitwood, Susan M. Rundell, Darren Y. Li, Quaneisha L. Woodford, Tommy T. Yu, Jose R. Lopez, Danny Greenblatt, Julie Kang, Jason P. Londo

**Affiliations:** Donald Danforth Plant Science Center, St. Louis, MO 63132 USA; Northern Iowa University, Department of Biology, Cedar Falls, IA 50614 USA; United States Department of Agriculture, Agriculture Research Service, Grape Genetics Research Unit, Geneva, NY 14456 USA

**Keywords:** Leaf morphology, leaf shape, phenotype, development, grape, climate, plasticity

## Abstract

The shape of leaves are dynamic, changing over evolutionary time between species, within a single plant producing different shaped leaves at successive nodes, during the development of a single leaf as it allometrically expands, and in response to the environment. Notably, strong correlations between the dissection and size of leaves with temperature and precipitation exist in both the paleorecord and extant populations. Yet, a morphometric model integrating evolutionary, developmental, and environmental effects on leaf shape is lacking. Here, we continue a morphometric analysis of >5,500 leaves representing 270 grapevines of multiple *Vitis* species between two growing seasons. Leaves are paired one-to-one, vine-to-vine accounting for developmental context, between growing seasons. Linear Discriminant Analysis reveals shape features that specifically define growing season, regardless of species or developmental context. The shape feature, a more pronounced distal sinus, is associated with the colder, drier growing season, consistent with patterns observed in the paleorecord. We discuss the implications of such plasticity in a long-lived woody perennial, such as grapevine, with respect to the evolution and functionality of plant morphology and changes in climate.

## Introduction

Despite the astonishing array of leaf shapes found in flowering plants, the underlying causes of such diversity—genetic and functional—remain largely a mystery. Leaf shape might contribute to any number of functions—including biomechanical support, light interception through the canopy, thermoregulation, regulating the boundary layer, hydraulic constraints, adaptations against herbivory, or developmental constraint (Givnish, 1979, 1987; Sack and Holbrook, 2006; Nicotra et al., 2011)—but it is also possible that aspects of shape are functionally neutral and vary across evolution through mechanisms other than natural selection. Known genetic, developmental, and environmental effects (described below) regulate leaf morphology. Yet, a comprehensive model integrating these factors—the development of differently shaped leaves produced by individual plants belonging to distinct species in varying real-world environments— remains rarely described. Such a comprehensive model is required to fully understand the evolution of plant morphology (Kaplan, 2001).

Genetically, molecular pathways with conserved function across the angiosperms alter conspicuous attributes of leaves, such as the role of KNOTTED1-LIKE HOMEOBOX (KNOXs) genes in promoting leaf complexity (Janssen et al., 1998; Bharathan et al., 2002; Kimura et al., 2008) and the activity of CUP-SHAPED COTYLEDON family members (CUCs) that alter leaf lobing and serration (Blein et al., 2008; Kawamura et al., 2010). From a quantitative genetic perspective, leaf shape is highly heritable, regulated by small effect loci that mostly remain uncharacterized at the molecular level (Langlade et al., 2005; Tian et al., 2011; Chitwood et al., 2013; Chitwood et al., 2014a).

Developmentally, the shape of a leaf is constantly changing throughout the course of development as it expands during ontogeny (which we define as the development of a single leaf, contrasting with heteroblasty described below). This was an early observation, made in the 1700s, when Stephen Hales punched holes in fig leaves and observed that different regions of the leaf expand at unequal rates (Hales, 1727). The patterns of allometric expansion in leaves vary between species (Das Gupta and Nath, 2015), regulated by cell proliferation and subsequent expansion (Avery, 1933; Poethig and Szymkowiak, 1995; Kang and Dengler, 2002). Beyond the dynamic shape changes in a single leaf, plants produce different types of leaves at different nodes, reflecting the temporal development of the meristem from which they arise, a process known as heteroblasty (Goebel, 1900; Ashby, 1948; Poethig 1990; 2010; Jones, 1993; Kerstetter and Poethig, 1998; Diggle, 2002). Changes in the timing of the heteroblastic progression of leaf shape can create evolutionary differences between species, a process known as heterochrony (Chitwood et al., 2012, 2014b; Cartolano et al., 2015).

Environment further modulates the complex genetic-developmental expression of leaf shape. A classic hypothesis proposes that the environment regulates leaf shape through timing and heteroblastic effects, explaining why juvenile-appearing leaves persist under low light conditions (Goebel, 1908; Allsopp, 1954). Careful morphological analysis of initiating leaves, however, suggests that such an interpretation is incorrect, and that the shape changes induced by environment (low light intensity) appear after leaf initiation. Thus the timing of the types of leaves a plant displays is unaffected (Jones, 1995). This interpretation is also supported by recent work analyzing the transcriptomic responses in leaf primordia to simulated foliar shade and heteroblasty, showing that these phenomena in tomato leaf primordia are largely distinct at the molecular level (Chitwood et al., 2015a). Environmentally induced changes in leaf shape through timing-dependent (heterochronic) or timing-independent mechanisms is important, since field-based observations demonstrate that leaves plastically respond to their climate (Royer et al., 2009). This plasticity often results in changes to marginal serrations and lobes which are morphological features explicitly modulated by heteroblastic pathways at the molecular level in the Brassicaceae (Rubio-Somoza et al., 2014), tomato (Chitwood et al., 2015a), and Proteaceae (Ostria-Gallardo et al., 2015). Extant species distributions and the paleobotanical record attest to changes in these features during the evolution of flowering plants, especially long-lived woody perennials (Bailey and Sinnott, 1915, 1916; Webb, 1968; Wolfe, 1978, 1979, 1993, 1995; Givnish, 1979, 1984; Hall and Swaine, 1981; Richards, 1996; Wilf, 1997; Wilf et al., 1998; Jacobs, 1999, 2002; Field et al., 2005; Traiser et al., 2005; Royer and Wilf, 2006; Peppe et al., 2011). Yet, field and paleobotanical measurements of leaf shape rarely account for developmental effects (neither ontogeny nor heteroblasty).

Here, we analyze shape differences in >5,500 grapevine leaves and match leaves, from the same vines and same developmental stages, between the 2012-2013 (Chitwood et al., 2015b) and 2014-2015 growing seasons. The depth of the distal sinus, specifically, is a shape feature that has been altered in all *Vitis* species analyzed across different developmental contexts. Our results show that environmental plasticity in leaf shape affects leaves independently from other evolutionary and developmental effects, and we discuss the implications of our findings for the interpretation of the paleobotanical record and the effect that future climate change will have on plant development, especially in long-lived woody perennials such as grapevine.

## Materials and Methods

### Germplasm, sample collection, and scanning

More than 5,500 grapevine leaves, representing 270 vines from 12 *Vitis* species, 4 *V. vinifera* hybrids, and 3 species from the related genus *Ampelopsis,* were sampled from the USDA germplasm repository in Geneva, NY. This collection was first sampled during the 2012-2013 growing season, and the methods and results from that analysis--which demonstrates that latent genetic and developmental shapes within a single leaf can be resolved--are described in Chitwood et al. (2015b). The population was similarly analyzed for the 2014-2015 growing season, and a combined comparative analysis with the 2012-2013 data is presented in the current study.

For each growing season, leaves were collected the second week of June. All leaves > ~1 cm in length, arising from a single shoot per vine, were analyzed. Leaves were sampled keeping track of the node from which they originated (beginning with the shoot tip and counting sequentially towards the shoot base), stacked in order, and sealed in a Ziploc bag, for which an air hole had been cut in the corner. Leaves were kept in a cooler in the vineyard and then subsequently scanned within 1-2 days.

Leaves were arranged on a scanner (Epson Workforce DS-50000, Suwa, Japan), in their order collected from the shoot, with small labels indicating node number. The abaxial side of the leaf was imaged and a ruler was included in the scan. Files were named by vine ID with an appended letter indicating the file position in the image series.

Climate data was retrieved from the Northeast Weather Association (NEWA) website via Cornell University (newa.cornell.edu, retrieved August 1, 2015). Daily summaries of minimum, average, and maximum temperatures, leaf wetness hours, and precipitation were analyzed for the Geneva, NY station at the Vineyard North Farm approximately three miles from the USDA germplasm location. For each of the growing seasons, data was analyzed starting March the previous year until the end of June.

### Landmarking

Previously (Chitwood et al., 2015b), we had used a series of 17 landmarks to morphometrically analyze leaves. In this work, we use a set of 21 landmarks, which have been applied to both 2012-2013 and 2014-2015 growing season data. Landmarks were placed by hand using ImageJ (Abramoff et al., 2004). Landmarks were placed on either the left or right side of the leaf, but we describe the order landmarks were placed using a left-side orientation: 1) left side of the proximal vein base, 2) right side of the proximal vein base/left side of the distal vein base, 3) right side of the distal vein base/left side of the midvein base, 4) right side of the midvein base, 5) distal base of petiolar vein, 6) proximal base of petiolar vein, 7) width of proximal vein at petiolar vein branch point, 8) distal base of distal vein branch, 9) proximal base of distal vein branch, 10) width of distal vein at branch point, 11) distal base of midvein branch, 12) proximal base of midvein branch, 13) width of midvein at branch point, 14) tip of petiolar vein, 15) tip of proximal lobe, 16) proximal sinus, 17) tip of distal vein branch, 18) tip of distal lobe, 19) distal sinus, 20) tip of midvein branch, 21) tip of the leaf (see **Fig. 1**). Using ggplot2 (Wickham, 2009) in R (R Core Team, 2014), graphs for landmarks from each image were visually checked for errors. If errors were detected, the landmarking was redone for those particular samples.

**Figure 1:**
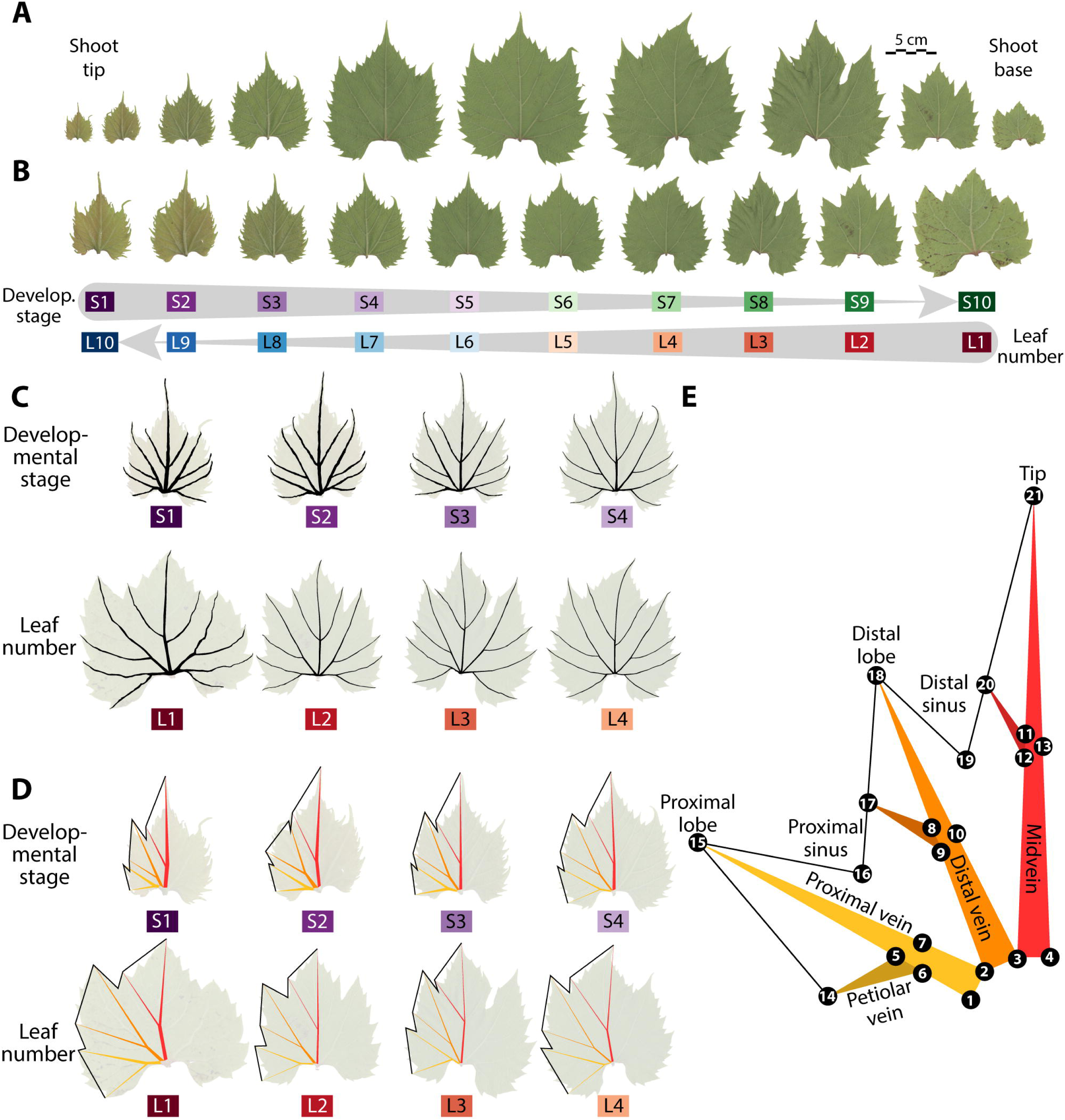
Developmental changes in leaf shape and landmarks. **A)** Leaves sampled along a shoot, from tip-to-base. **B)** Same leaves in A) except scaled to leaf length. The reciprocal relationship between developmental stage (Sn), sensitive to leaf ontogeny, and leaf number (Ln), sensitive to heteroblastic changes in leaf shape, is shown. **C)** From the first four developmental stages (S1-S4) and leaf numbers (L1-L4) shown for the shoot in A-B), the thickness of veins are highlighted. Note the dramatic decrease in vein thickness through progressive developmental stages (Sn) and leaf numbers (Ln). **D)** Shape information captured in leaves using landmarks. **E)** 21 landmarks used in this study capture leaf shape and vein patterning and thickness. Red indicates midvein, orange distal vein, yellow proximal vein. Darker shades indicate branching veins. Numbers indicate order landmarks were placed. Developmental stage indicated by purple (S1) to green (S10) and leaf number indicated by red (L1) to blue (L10) transitions.

### Morphometric analysis, statistics, and visualization

A Generalized Procrustes Analysis (GPA) was performed to allow super-imposed coordinates from leaves to be analyzed. The procGPA function (with reflect=TRUE) from the R package shapes (Dryden, 2013) was used. Eigenleaves were visualized using the shapepca function. PC scores, percent variance explained by each PC, and Procrustes-adjusted coordinates were obtained from the procGPA ouput.

For analyses determining Linear Discriminants defining growing season, species, or developmental shape attributes, or other pair-wise comparisons between growing seasons, a matched dataset between growing seasons was created. Leaves were matched one-to-one between growing seasons based on 1) originating from the same vine and 2) arising from the same shoot position. To match by shoot position, two matched datasets were created that were analyzed in parallel or appropriate for the analysis at hand: one dataset matched by vine and developmental stage (Sn) (**Dataset S1**) and another dataset matched by vine and leaf number (Ln) (**Dataset S2**). These datasets resulted in a total number of 2,243 leaf pairs matched by developmental stage (Sn) representing 223 vines and 2,245 leaf pairs matched by leaf number (Ln) representing 223 vines between growing seasons. Because ontogenetic and heteroblastic effects are strongest at the shoot tip and base, respectively (Chitwood et al., 2015b), only S1-S10 and L1-L10 were subsequently analyzed for each dataset, reducing the effective number of leaf pairs ultimately used in this study to 2,055 and 2,054 for developmental stage (Sn) and leaf number (Ln) datasets, respectively.

Linear Discriminant Analysis (LDA) on Procrustes-adjusted coordinates on matched datasets was performed using the lda function from the MASS package (Venables and Ripley, 2002). Growing season, developmental stage, leaf number, and species were all analyzed independent of each other. The predict function (stats package) and table function (base package) were used (dependent on MASS) to reallocate leaves (whether by growing season, developmental stage, leaf number, or species) using the linear discriminants. For LDAs performed by developmental stage (Sn) or leaf number (Ln), only the first 10 nodes in the series were considered. Results from the eight most abundant species (*Vitis acerifolia, V. aestivalis, V. amurensis, V. cinerea, V. labrusca, V. riparia, V. rupestris, V. vulpina*) were selected for visualization purposes.

Visualization was performed in ggplot2 (Wickham, 2009) using geom_area, geom_boxplot, geom_histogram, geom_point, geom_segment, and stat_smooth functions, among others, and color schemes derived from colorbrewer2.org.

## Results and Discussion

### Allometry

In a previous study, we had measured leaves from the 2012-2013 growing season with 17 landmarks describing the lobes, sinuses, and major vein branching points on both sides of the leaf (Chitwood et al., 2015b). Unlike in animal studies, allometry—changes in shape that vary by organ size, or the growth of body parts at different rates—is not a straightforward concept when applied to plants because of their iterative growth. For example, leaves are smaller in overall size at the base *and* tip of the growing shoot (**Fig. 1A**). The older, first-formed leaves at the base of the shoot are mature and have reached their developmental plateau, ceasing to expand, and are smaller in size because they are genetically specified to be so. The young leaves at the tip of the shoot are smaller because they are the youngest leaves and have not completed blade expansion to reach maturity.

Because small leaves are associated with two different phenomena—the different shapes and sizes of leaves at successive nodes (heteroblasty) and the development of individual leaves as they expand (ontogeny)—an allometric analysis that accounts for shape differences attributable to size would only confound these distinct effects (Chitwood et al., 2015b). To circumvent this, leaves were cataloged by two, complementary indices. Developmental stage (Sn) counts nodes starting at the shoot tip to the shoot base (S1 starting at the shoot tip). Leaf number (Ln) counts nodes starting at the shoot base to the shoot tip (L1 starting at the shoot base) (**Fig. 1B**). Developmental stage (Sn) is sensitive to ontogenetic effects attributable to the expansion of a single leaf as it grows, as leaves at the tip are young and rapidly increasing in size. Leaf number (Ln) is sensitive to heteroblastic effects, as leaves at the base of the shoot are the oldest in the leaf series, and comparing shapes between them represents intrinsic shape differences between leaves at maturity. Even though developmental stage (Sn) and leaf number (Ln) are inversely related, their shape effects are separable and reflect distinct phenomena, as we have previously shown (Chitwood et al., 2015b).

An attribute of leaves our previous landmarks failed to measure was vein thickness. When leaves were scaled to size from the shoot tip onwards (Sn), the leaves showed a visibly sharp decrease in the overall area of the major veins relative to blade regions (**Fig. 1C**). This was also true when looking at leaves originating from the shoot base onwards (Ln). For this study, we reanalyzed the leaves from the 2012-2013 growing season and analyzed for the first time 2014-2015 leaves using a new set of 21 landmarks on only one side of the leaf that measured the thickness of veins (**Fig. 1D-E**). Unlike our previous landmarks, our new landmarks capture an effect related to vein thickness that affects smaller leaves at the shoot tip and base alike (**Fig. 1**), allowing a traditional allometric analysis based on leaf size alone to be performed. Plotting the natural log of each of the vein areas (**Fig. 2A**) or blade areas (**Fig. 2B**) against the natural log of the overall leaf area reveals near-linear relationships between sub-regions of the leaf with overall leaf size. If these allometric relationships are plotted against each other, it becomes apparent that blade regions expand faster than vein as leaf size increases (**Fig. 2C**). These relationships are strongly linear in the natural log-transformed space and represent leaves from different species, developmental stages throughout the shoot, and different years of collection, showing that the size of different leaf sub-regions—whether vein or blade—relative to leaf size is a strongly conserved feature of grapevine leaves.

**Figure 2:**
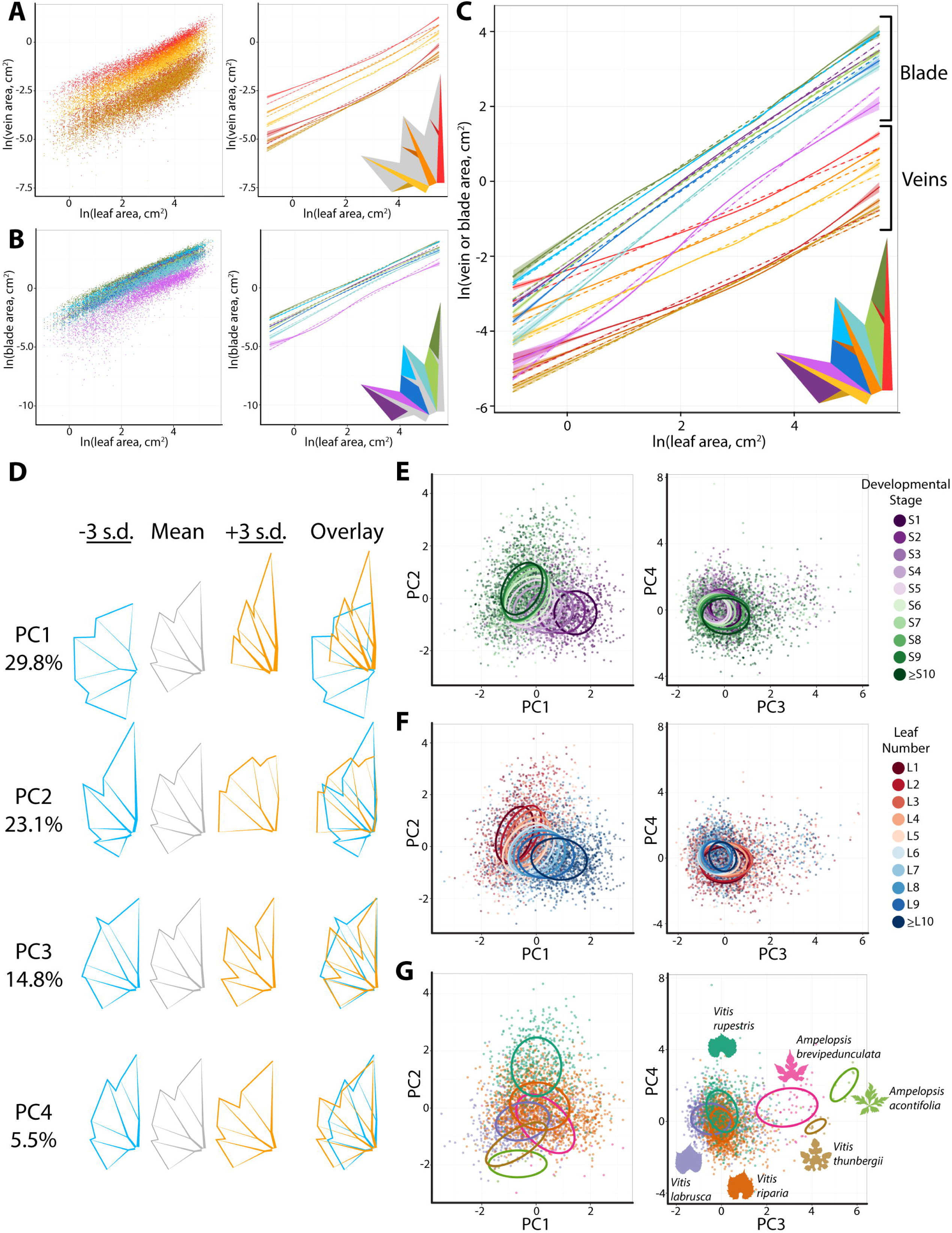
Allometry and morphospace. **A)** Natural log of the indicated vein areas by color plotted against natural log of overall leaf area. Left, scatterplot. Right, both linear (solid line) and non-linear local regression (dotted line) curves with 95% confidence bands. **B)** Same as A) except plotting the natural log of the indicated blade areas by color against the natural log of overall leaf area. **C)** As in A-B), plots of the natural log of vein and blade areas against the natural log of overall leaf area indicated by color. Note that blade areas increase more rapidly than vein as leaves become larger. **D)** Eigenleaves representing the overall morphospace of >5,500 leaves analyzed from all species, developmental contexts, and growing seasons. Shown are representative leaves at −/+ 3 standard deviations (“s.d.”; indicated by blue and orange, respectively) and overlay. Leaves are aligned at the petiolar junction for ease in comparing. **E)** Developmental stage, **F)** leaf number, and **G)** species identity projected onto the morphospace. For each level of a factor, 95% confidence ellipses are drawn. Only a subset of morphologically diverse species, and the first 10 nodes of developmental stage and leaf number, is shown for visual clarity. Species indicated by various colors. Developmental stage indicated by purple (S1) to green (S10) and leaf number indicated by red (L1) to blue (L10) transitions.

### Morphospace

A Principal Component Analysis (PCA) performed on Procrustes-adjusted coordinates (performed with scaling and reflection) reveals the genetic and developmental context of leaf shape data and principal sources of shape variance (**Fig. 2D**). The first PC accounts for 29.8% of shape variance and is defined by shape features related to the length-to-width ratio of leaves, the length of the midvein, the amount of blade outgrowth at the leaf base, the angles of the major veins, and importantly the thickness of the veins (**Fig. 2D**). Both developmental stage (Sn) and leaf number (Ln) mostly vary by PC1. Young leaves at the shoot tip (low Sn values) are associated with high PC1 values (**Fig. 2E**) and thick veins, long midveins, and minimal laminar outgrowth at the leaf base. Mature leaves at the shoot base (low Ln values) are associated with low PC1 values (**Fig. 2F**) and wider leaves with extensive blade outgrowth at the leaf base.

Different *Vitis* species vary mostly by PC2 and PC3 (**Fig. 2G**). PC2 (accounting for 23.1% of shape variance) explains variation related to length of the midvein and outgrowth at the leaf base, but such that a longer midvein and outgrowth at the leaf base occur together (rather than the opposite configuration seen in PC1). *Vitis rupestris* has exceptionally high PC2 values relative to other species, as is expected considering the characteristic short midvein and lack of blade outgrowth at the base of its leaves. PC3 (accounting for 14.8% of the shape variance) explains variation related explicitly to lobing (**Fig. 2D**). High PC3 values separate deeply lobed species (such as *V. thunbergii, A. brevipedunculata,* and *A. acontifolia)* from less lobed species (such as *V. riparia, V. labrusca,* and *V. rupestris)* (**Fig. 2G**).

We note that, because PCs are orthogonal with each other, the restriction of most variance associated with shoot position to PC1 (**Fig. 2E-F**) and most variance associated with species identity to PCs 2 and 3 (**Fig. 2G**) suggests that shape differences due to genetic and developmental effects are largely independent from each other. This idea, that a single leaf is a composite of latent genetic and developmental shapes (i.e., multivariate signatures that differentially define genetic vs. developmental shape variance), and that such latent shapes can be resolved, is detailed in our previous work analyzing only the 2012-2013 data with 17 landmarks (Chitwood et al., 2015b).

### Interannual variability

Genetic and developmental effects substantially affect the leaf morphospace (**Fig. 2D-G**). We must account for these strong effects if leaf shape differences between growing seasons are to be examined. In our dataset, leaves were harvested from the same vines for the 2012-2013 and 2014-2015 growing seasons. We decided to create paired datasets based on our two indexing methods—developmental stage (Sn) and leaf number (Ln) (**Fig. 1A-B**). Leaves were paired between the 2012-2013 and 2014-2015 growing seasons, one-to-one, originating from 1) the same vines and 2) corresponding to one of two different developmental contexts, developmental stage (Sn) or leaf number (Ln).

A Linear Discriminant Analysis (LDA) performed on the paired data identifies the linear combination of Procrustes-adjusted coordinate landmarks most discriminating between growing seasons. Distributions of Linear Discriminant 1 (LD1) values are visibly separated for leaves from the 2012-2013 and 2014-2015 growing seasons (**Fig. 3A**). Importantly, there are no strong biases in the ability of LD1 values to discriminate growing seasons across the sampled developmental stages (Sn) and leaf numbers (Ln) (**Fig. 3B**) nor species (**Fig. 3C**), demonstrating that the shape features most discriminating different years affect leaves from all genetic and developmental contexts relatively equally. Using a discriminant space trained on year (without regard to species or developmental context) approximately two-thirds of leaves can be correctly classified to the correct year (**Table 1**). We conclude that the shape attributes influenced by interannual variability can be used to predict leaf identity independently of genetic and developmental effects.

**Table 1:** Prediction of leaf growing season.

| Index | Actual year | Predicted correct | Predicted wrong | Percent correct |
| --- | --- | --- | --- | --- |
| Dev. stage (Sn) | 2013 | 1346 | 709 | 65.5% |
| Dev. stage (Sn) | 2015 | 1431 | 624 | 69.6% |
| Leaf number (Ln) | 2013 | 1393 | 661 | 67.8% |
| Leaf number (Ln) | 2015 | 1420 | 634 | 69.1% |

**Figure 3:**
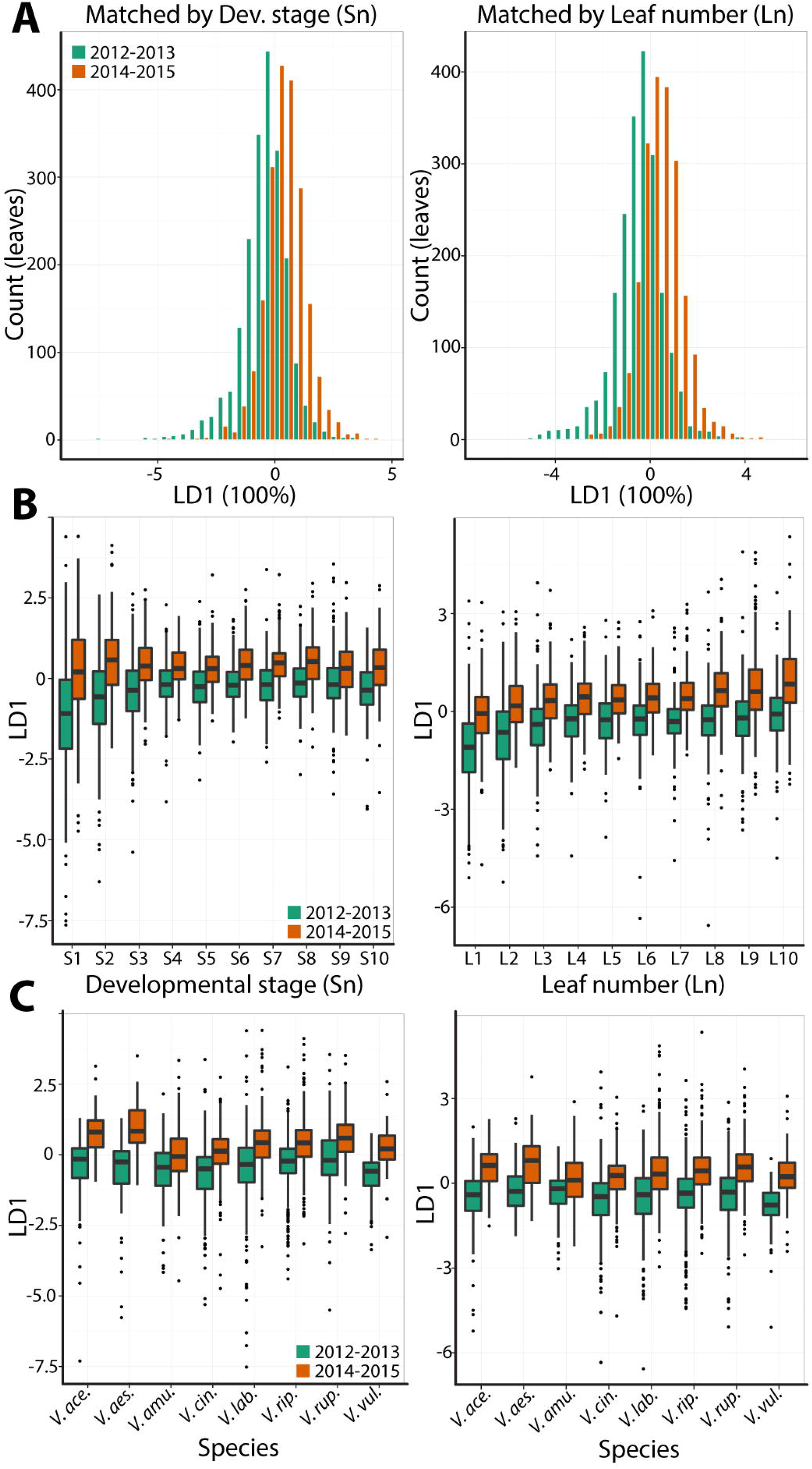
Linear discriminants for growing season. **A)** Histograms for linear discriminant values of leaves collected in different growing seasons (green, 2012-2013; orange, 2014-2015) for datasets matched by developmental stage (Sn; left) and leaf number (Ln; right). **B)** For matched datasets by developmental stage (left) and leaf number (right) linear discriminants similarly discriminate growing seasons regardless of developmental context. **C)** Similar to B), datasets matched by developmental stage (left) and leaf number (right) discriminate between growing seasons similarly across species. Only the eight most abundantly represented species in the dataset are shown for visual clarity. *V. ace.=Vitis acerifolia*; *V. aes.=V. aestivalis; V. amu.=V. amurensis; V. cin.=V. cinerea, V. lab.=V. labrusca; V. rip.=V. riparia; V. rup.=V. rupestris; V. vul.=V. vulpina.*

Which shape features most discriminate leaves between years? By visualizing the mean shapes of leaves predicted as arising from the 2012-2013 and 2014-2015 growing seasons, it becomes evident that the distal sinus in leaves from the 2014-2015 season is deeper than the distal sinus in leaves from the 2012-2013 growing season (**Fig. 4A**, arrow). But does this shape feature—a pronounced distal sinus—coincide with shape features associated with developmental context or species?

**Figure 4:**
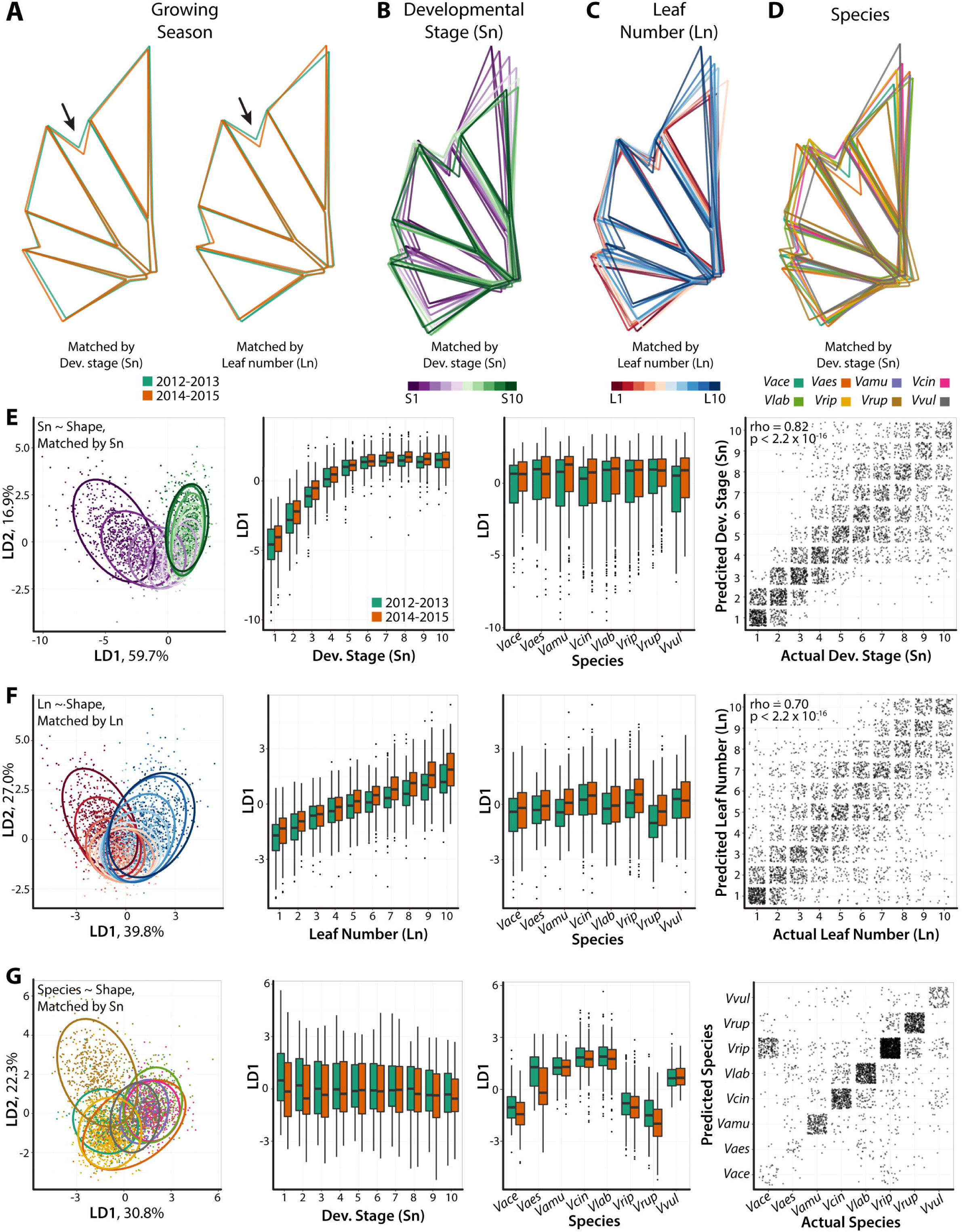
Shape features discriminating factors and their independence from each other. **A)** Mean shapes from leaves predicted arising from the 2012-2013 (green) and 2014-2015 (orange) growing seasons using linear discriminants (see **Fig. 3**). Both datasets matched by developmental stage (Sn; left) and leaf number (Ln; right) show a more pronounced distal sinus (indicated by arrow) in the 2014-2015 growing season compared to 2012-2013. **B-C**) Similar to A) except showing mean leaves resulting from a discriminant analyses performed by B) developmental stage (Sn; S1, purple to S10, green) or C) leaf number (Ln; L1, red to L10, blue). Leaf length and blade outgrowth at the leaf base are the most affected shape features. **D)** Similar to A-C) except showing mean leaves resulting from a discriminant analysis performed by species. Only the mean leaves of the eight most abundant species are shown, indicated by color, for clarity. **E-G)** Results from discriminant analyses performed by E) developmental stage (Sn), F) leaf number (Ln), and G) species. Discriminant values separate by the factor for which they are specified, but not others (including growing season), demonstrating independence of discriminant shape features for different factors. Far left: discriminant space based on LD1-2 values for each respective analysis. 95% confidence ellipses are provided for factor levels, indicated by colors. Middle left: LD1 values from respective analyses across developmental contexts (developmental stage, Sn, or leaf number, Ln, as appropriate for the analysis), separated by growing seasons. Middle right: LD1 values from respective analyses across the eight most abundant species, separated by growing seasons. Far right: confusion matrices for predicted factor level against actual factor level. Each point represents a single leaf. Spearman’s rho and p-values provided for correlation between predicted and actual identities in the developmental stage and leaf number datasets. *Vace=Vitis acerifolia; Vaes=V. aestivalis; Vamu=V. amurensis; Vcin=V. cinerea, Vlab=V. labrusca; Vrip=V. riparia; Vrup=V. rupestris; Vvul=V. vulpina.*

To answer these questions, we performed similar Linear Discriminant Analyses (LDAs) on developmental stage (Sn), leaf number (Ln), and species identity, as the sole factors to be discriminated without regard to other factors. The shape features defining developmental stage (Sn) (**Fig. 4B**) and leaf number (**Fig. 4C**) both concern the length of the midvein and the amount of blade outgrowth at the leaf base. The discriminant analysis for species reveals that leaves arising from different species are defined by a number of features in different combinations, mostly related to the length of the midvein, length-to-width ratio, and in some species (especially *V. aestivalis*) the depth of the sinus (**Fig. 4D**). These features are largely distinct from the features that define growing season. If LD1 values from developmental stage (Sn, **Fig. 4E**), leaf number (Ln, **Fig. 4F**), and species (**Fig. 4G**) are compared across developmental contexts and species, the linear discriminant correctly differentiates the factor it is designed to discern, but not growing season. Confusion matrices (**Fig. 4E-G**) demonstrate the predictive power of these features for each factor, but each discriminant analysis is specific to its intended factor and distinct from growing season.

Regardless of developmental stage (Sn) or leaf number (Ln) (**Fig. 3B**), or species (**Fig. 3C**), the depth of the distal sinus is a shape feature discriminating leaves from different growing seasons. The distal sinus (**Fig. 4A**) is an attribute of grapevine leaves that changes between years on the same vines, largely independent from other shape features specific to developmental stage (**Fig. 4B, 4E**), leaf number (**Fig. 4C, 4F**), or species (**Fig. 4D, 4G**).

### Implications for the paleorecord and-future climate change

Only two factors vary between the 2012-2013 and 2014-2015 growing seasons: 1) climate and 2) the age of the vines. We assume that two years, in the overall lifespan of a long-lived woody perennial plant like grapevine, negligibly affects leaf shape compared to changes in climate. Climate can potentially influence leaf development in grapevine over a two-year period, as leaves are patterned within buds the year prior to the growing season in which they emerge (Carmona et al., 2008). For this reason, we evaluated weather data starting with the years prior to collection (2012 and 2014) when temperatures began to rise above freezing (around March) and through collection time (mid June) to the end of June in the subsequent years (2013 and 2015).

Temperatures were colder during the 2014-2015 growing season compared to 2012-2013 (**Fig. 5A**). The pattern of cumulative precipitation is more complicated: 2014 had more precipitation than 2012, but 2013 had more precipitation than 2015 (**Fig. 5B**). If instead leaf wetness hours are measured (which is influenced not only by precipitation, but solar radiation, wind, humidity, etc.) then the 2014-2015 growing season appears to be drier than 2012-2013 (**Fig. 5C**). Overall, 2014-2015 appears to be colder and drier (at least by leaf wetness hours) than 2012-2013 and produces leaves with deeper distal sinuses regardless of developmental context or species identity (**Figs. 3-4**).

**Figure 5:**
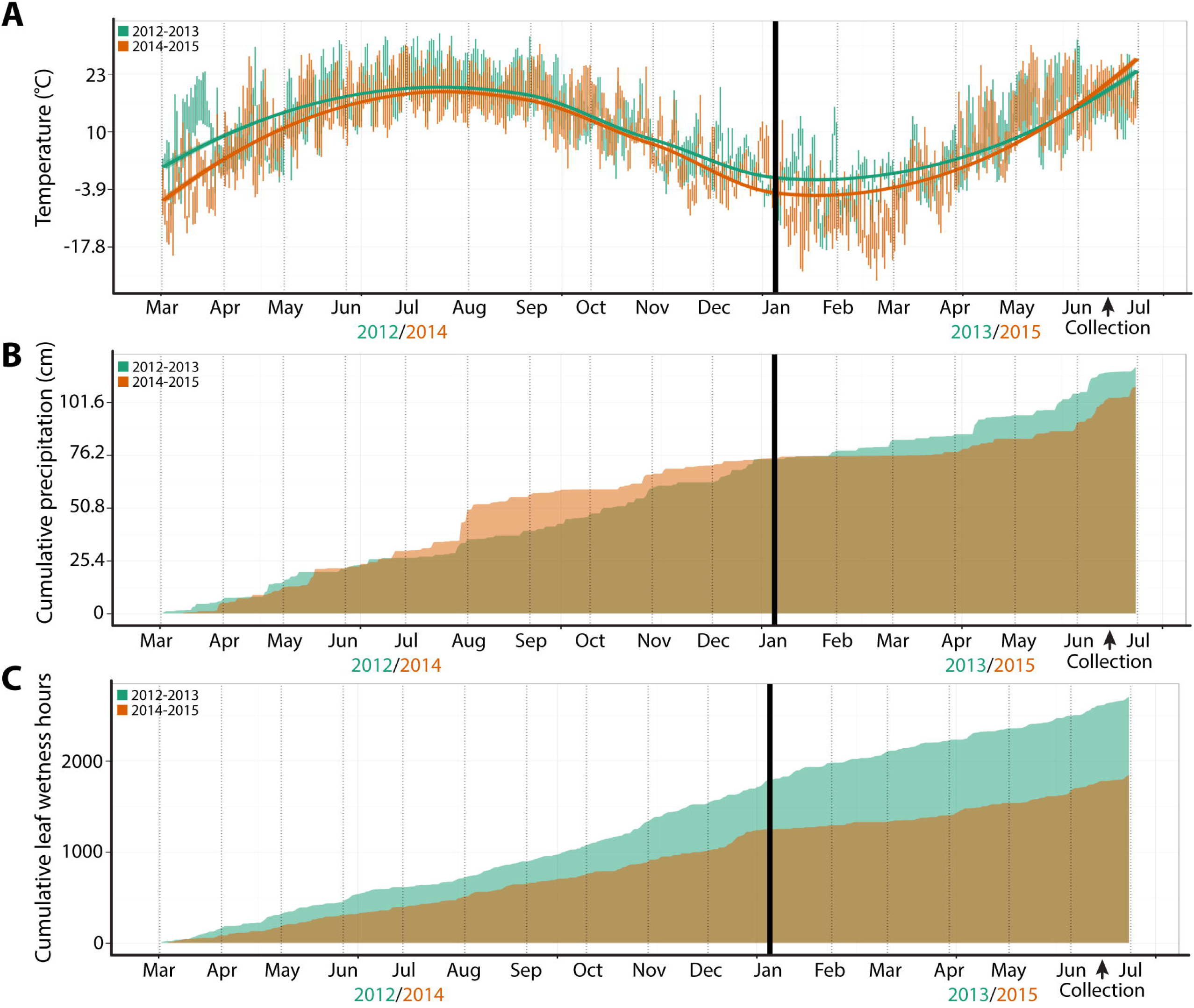
Climate data for the 2012-2013 and 2014-2015 growing seasons. **A)** Daily temperatures for the 2012-2013 (green) and 2014-2015 (orange) growing seasons. Bars indicate minimum and maximum daily temperatures fitted with a local regression curve for average daily temperature (95% confidence bands indicated). **B)** Cumulative precipitation (cm) and **C)** cumulative leaf wetness hours between growing seasons. Colors are transparent, and overlap in cumulative plots is brown.

We stress that our data measures, in a manner accounting for both species diversity and plant development, interannual variability in grapevine leaf shape between two years. Additional years must be analyzed to truly correlate leaf shape features with climate. But the observation of a pronounced distal sinus in the colder, drier year is of interest given the well-established connection between leaf dissection and the paleoclimate. Multiple studies dating back a century (Bailey and Sinnott, 1915) observe increased leaf serration depth—in both extant and fossil populations—in colder and drier climates (e.g., temperate) compared to those that are warmer and wetter (e.g., sub-tropical and tropical) (Bailey and Sinnott, 1916; Webb, 1968; Wolfe, 1978, 1979, 1993, 1995; Givnish, 1979, 1984; Hall and Swaine, 1981; Richards, 1996; Wilf, 1997; Wilf et al., 1998; Jacobs, 1999, 2002; Field et al., 2005; Traiser et al., 2005; Royer and Wilf, 2006; Peppe et al., 2011). Although the functional significance of such correlations remains a point of contention, and might even reflect strong developmental constraints, the role of the environment in shaping leaf morphology over evolutionary time cannot be denied.

This study, however, specifically looks at plasticity. Transplant experiments in *Acer rubrum* demonstrate both genetic and plastic environmental effects consistent with a more dissected leaf morphology in temperate compared to tropical environments (Royer et al., 2009). This study, however, examined the same individuals between years, analyzing the shape differences due to interannual climate variability in the same vines. The main shape attribute that altered was the depth of the distal sinus, regardless of developmental context or species. Leaf lobing, specifically, is a feature that varies plastically within the canopy of a single plant (Gray, 1990; Zwieniecki et al., 2004; Sack et al., 2006), by nutrient composition and availability (Gosler et al., 1994; Dorken and Barrett, 2004), planting density (Semchenko and Zobel, 2007), and natural versus controlled environments (McLellan, 2000), light intensity (Jones, 1993, 1995) and can vary as much as by environmental effects as genetic (Gosler et al., 1994). Such sensitivity in a morphological feature reflects either adaptive significance or a developmental constraint resulting from the plastic response of a plant to differing environmental circumstances.

Grapevine, in particular, is a focus of climate and agricultural productivity models to determine the disruptive influence of climate change on crop outputs (Jackson and Cherry, 1988; Kenny and Harrison, 1993; Bindi et al., 1996a, 1996b; Schultz et al., 1998; Schultz, 2000; Kolb et al., 2001; Jones et al., 2005; White et al., 2006; Moutinho-Pereira et al., 2009; Rogiers et al., 2011). The ultimate extent that climate change will disrupt the current biogeographical distribution of wine-making is contentious (Hannah et al., 2013), in part because of adaptations in viticultural practices to accommodate a warmer climate (i.e., plasticity) (van Leeuwen et al., 2013). Long-lived woody perennials, such as grapevine, provide a unique opportunity to study the role of climate in altering plant development, as their longevity allows for sizeable sampling of changing climate conditions for the same individual in the same geographic location. The unique leaves of *Vitis* spp. allow homologous features to be tracked, a property not necessarily afforded in the shapes of leaves from other species. Our results demonstrate that genetic, developmental (ontogenetic and heteroblastic), and environmental effects influence leaf shape in a manner largely independent of one another and can be resolved as latent shape features within the composite shape of a single leaf (Chitwood et al., 2015b). Changes in climate affect the morphology and developmental patterning of grapevine leaves in specific ways that universally affect all developmental stages across species within the genus *Vitis.* The morphological responses of plants to climate not only serve as bio-statistical indicators of the impact of a changing environment on how plants develop, but can potentially reveal insights into the distinction between adaptive versus constrained morphological features that vary over geological time.

---

**Dataset S1: Procrustes-adjusted coordinates for leaves matched by vine and developmental stage (Sn) between growing seasons.** A table providing growing season (“year”), file name of original scan (“label”), vine identification (“vine”), the pixels per cm^2^ for the scan (“px2_cm2”), species (“species”), the number of leaves per shoot (“max_count”), the developmental stage (Sn, “leaf_stage”), the leaf number (Ln, “leaf_num”), and Procrustes-adjusted coordinates (x1, y1 to x21, y21) for leaves matched by vine and developmental stage between growing seasons.

**Dataset S2: Procrustes-adjusted coordinates for leaves matched by vine and leaf number (Ln) between growing seasons.** A table providing growing season (“year”), file name of original scan (“label”), vine identification (“vine”), the pixels per cm^2^ for the scan (“px2_cm2”), species (“species”), the number of leaves per shoot (“max_count”), the developmental stage (Sn, “leaf_stage”), the leaf number (Ln, “leaf_num”), and Procrustes-adjusted coordinates (x1, y1 to x21, y21) for leaves matched by vine and leaf number between growing seasons.

